# Salicyl-Carnosine Protects Primary Cortical Rat Neuron Cultures in Conditions of Oxygen-Glucose Deprivation and NMDA-Induced Excitotoxicity by Preventing Oxidative Stress

**DOI:** 10.64898/2026.08.07.743511

**Authors:** Alexander V. Lopachev, Rogneda B. Kazanskaya, Olga I. Kulikova, Anastasiya V. Khutorova, Denis A. Abaimov, Tatiana N. Fedorova

## Abstract

Therapy of ischemic stroke is currently limited to pharmacological and/or mechanical recanalization. There are no neuroprotective therapies approved for use during the rehabilitative phase of ischemic stroke, which is characterized by neurodegenerative changes. Thus, the search for neuroprotective compounds capable of preventing neuronal death caused by pathogenetic cascades triggered during hypoxia is an urgent task. In this study, we demonstrate increased culture viability following pre- and post-incubation with salicyl-carnosine (SC) in a model of oxygen glucose deprivation on a primary culture of rat cortical neurons. Its neuroprotective properties were greater than that of acetylsalicylic acid and carnosine, and it was effective in lower concentrations. In addition, SC protected the culture from NMDA-induced excitotoxicity. We also showed the passage of SC into neurons, and the presence of its direct antioxidant activity in a model of paraquat-induced oxidative stress. The neuroprotective effects of SC are associated with a decrease in the level of pro-apoptotic protein Bak and a decrease in the activation of kinase p38, as well as an increase in the activation of kinase ERK1/2. The acquired data suggests that SC is a promising neuroprotective compound, and warrants further investigation in vivo.

## INTRODUCTION

Acute ischemic stroke is one of the most common cerebrovascular pathologies [1]. By the age of 80, the risk of ischemic stroke reaches approximately 25%, while the [2, 3] risk of ’silent’ stroke approaches 100% [4]. The time-course of ischemic stroke can be divided into two phases – the acute phase (minutes/hours/up to 21 days) and the chronic phase (days/weeks/months). The acute phase begins immediately following vessel thrombosis and is characterized by acute tissue hypoxia. During this phase, hypoxia induces neuron loss and initiates various pathogenic pathways. The oxidative stress, inflammation, and excitotoxicity which originate in this phase lead to neurodegeneration during the second, rehabilitative phase [5]. Following rehabilitation, the risk of recurrent ischemic stroke constitutes 2.5 – 4.0 % [2, 3].

Excitotoxicity is one of the most well-studied causes of neuron loss during the acute phase of ischemic insult. ATP deficiency disrupts ionic homeostasis, leading to neuronal depolarization, impaired transport processes, and increased release of various neurotransmitters, including glutamate. This accumulation of extracellular glutamate results in AMPA and NMDA receptor hyperactivation, causing sodium and calcium ions to enter the cytoplasm [6]. Calcium influx results in the generation of reactive oxygen species (ROS) and mitochondrial damage, which subsequently leads to neuron loss through necrosis or apoptosis. Oxidative stress (OS), induced by hypoxia and excitotoxicity, is one of the primary mechanisms of neuron loss during the rehabilitation phase [7]. Following reperfusion, ATP synthesis increases rapidly, causing an elevation in intracellular ROS [8]. This rise in ROS levels triggers the activation of pro-apoptotic and pro-necrotic signaling pathways, leading to further neuronal loss.

Glucose-oxygen deprivation (OGD), NMDA-induced excitotoxicity, and mitochondrial toxin models are used to model ischemia, excitotoxicity, and OS on neuronal cell cultures, respectively [9–11]. These models not only allow rapid, relative to animal models of cerebral stroke, screening of candidate compounds for neuroprotective activity, but also enable the identification of their mechanisms of action.

The aim of therapeutic intervention for ischemic stroke is to shorten the duration of the acute phase via intravenous administration of thrombolytic drugs such as the recombinant tissue-type plasminogen activator and/or mechanical recanalization [12]. However, there is currently no clinically approved therapy that targets neuron loss at the cellular level. Administering Aspirin (acetylsalicylic acid, ASA) to patients before or immediately after an ischemic stroke has been shown to reduce the amount of brain tissue damage and neurological symptoms [13, 14].

ASA inhibits the activity of cyclooxygenases 1 and 2 through irreversible binding [15]. As such, ASA blocks thromboxane A2, preventing platelet aggregation [16], which is why it is often used to prevent thrombosis [17]. A meta-analysis of studies on the effectiveness of low-dose ASA in the recovery phase following acute ischemic stroke has shown that if therapy is initiated within the first two days following ischemic stroke, it can improve the overall recovery prognosis and reduce the risk of recurrent ischemic stroke without the high risk of hemorrhagic complications [18]. However, while aspirin remains the undisputed gold standard of antiplatelet therapy, its use is complicated by side effects and aspirin resistance. Studies suggest that between 5% and 45% of patients have a reduced response to aspirin, likely due to differences in genetic background [19].

Unlike thrombolytics, neuroprotective compounds can affect the outcome of both the acute and rehabilitative phases of ischemic stroke. Neuroprotective drugs may prevent the initiation of pathogenic pathways during the acute phase and reduce their impact during the rehabilitative phase. The search for neuroprotective agents for the treatment of ischemic insult is crucial [20], particularly following the development of mechanical recanalization techniques. It was shown that ASA is neuroprotective in an in vitro model of brain ischaemia at 0.1-1 mM [21].

Carnosine (β-alanyl-L-histidine, CN) is an endogenous dipeptide found in excitable tissue, including muscle and nervous tissues. It possesses multiple properties that contribute to its neuroprotective efficacy, including direct and indirect antioxidant action, anti-glycation, metal-chelating, chaperone-like, and pH-buffering actions. The neuroprotective effects of CN have been demonstrated in both *in vitro* and *in vivo* models of brain injury [22]. In models of direct induction of OS on primary rat neuronal cultures by rotenone and AAPH, it reduces the amount of ROS in cells [23], and protects neurons from the excitotoxic effects of NMDA [24]. *In vivo*, CN has been shown to be effective in models of parkinsonism [25, 26] and ischemic stroke, including focal and global ischemia [27–29]. In a model of focal cerebral ischemia in the middle cerebral artery basin, CN prevented the activation of pro-apoptotic signaling cascades in the ischemic penumbra zone, thereby reducing the infarct volume [30]. Furthermore, neurons actively uptake CN via PEPT2 transporters [31]. However, the clinical applicability of CN is limited due to its rapid degradation by serum and tissue carnosinases [22].

Drugs with combined action, possessing both antiplatelet and neuroprotective properties, are of particular interest in the context of acute ischemic stroke and the prevention of recurrent ischemic stroke.

Salicyl-carnosine (SC) is a compound synthesized from CN and ASA. SC possesses significant antioxidant and antiplatelet effects. Its antiplatelet properties are comparable to those of ASA, as demonstrated in conditions of ADP-induced platelet aggregation. At the same time, SC does not cause gastric ulcers, a known side-effect of ASA [32]. This facilitates its long-term administration, such as for preventing recurrent ischemic stroke.

In the context of the aforementioned properties of SC, it is appropriate to study its effects in ischemic stroke models. One of the most common *in vitro* models of ischemic stroke is oxygen glucose deprivation (OGD) of neuronal cultures, which mimics the conditions of hypoxia observed in ischemic stroke [33]. This study assesses the neuroprotective effects of SC in rat cortical neuron cultures subjected to OGD, and in conditions of NMDA-induced excitotoxicity.

## MATHERIALS AND METHODS

### 2.1. Obtaining the primary culture of rat cortical neurons

Rat cortical tissue was obtained from 18-day-old Wistar rat embryos. The tissue was rinsed in Ca^2+^- and Mg^2+^-free Hanks’ solution (PanEco, Russia) and cleaned of vessels. The tissue was then incubated for 15 min at 37°C in trypsin-EDTA solution (PanEco, Russia).

Trypsin was inactivated by adding 20% fetal calf serum (FBS) (BioSera, UK). Tissue was rinsed twice with Hanks’ solution and suspended in MEM medium (PanEco, Russia) with 10% FBS and 100 units/mL penicillin-streptomycin (PanEco, Russia). The obtained suspension was centrifuged for 3 min at 300 g and resuspended in MEM medium (PanEco, Russia) with the above-mentioned additives. The obtained pure suspension was evenly distributed at a density of 1.2-105 cells per cm^2^ into 96-well plates (Nunc, USA), and at a density of 1.2^10^5^ cells per cm^2^ 24-well plates (SPL, Republic of Korea) which were pretreated with 0.1 mg/mL poly-L-ornithine for 12 h (Sigma, USA). Cultures were maintained in a CO_2_ incubator (SHEL LAB, USA) at 37°C, 90% humidity, 5% CO_2_ for 24 hours. The medium was then replaced with Neurobasal Medium (Gibco, USA) with 100 units/mL penicillin-streptomycin, 1% GlutaMAX (Gibco, USA) and 2% serum-free B-27 supplement (Gibco, USA). Cultures were maintained in a CO_2_ incubator for 12-14 days. Every two days, half of the medium volume was replaced with fresh medium.

### 2.2. Oxygen glucose deprivation

Brain ischemia was induced in a primary culture of rat cortical neurons by subjecting them to 3 hours of oxygen glucose deprivation (OGD) followed by 21 hours of reoxygenation. The duration of GCD was selected such that culture viability was reduced by ∼50%, guided by publicly available protocols [33, 34]. The results of the selection of OGD durations are presented in Figure 1S in the supplementary materials. The experiments were conducted on days 10-12 after culture seeding. Prior to OGD, the culture medium was replaced with glucose-free artificial cerebrospinal fluid (aCSF) containing 125 mM NaCl, 26 mM NaHCO_3_, 4 mM KCl, 1.25 mM NaH_2_PO_4_, 1.2 mM MgCl_2_ and 2 mM CaCl_2_. The cells in the control group were given aCSF with a final glucose concentration of 25 mM. To induce oxygen deprivation, plates from the hypoxia group were placed in a New Brunswick Galaxy 48 R hypoxic chamber (Eppendorf, Germany) for 4 hours at 1% O_2_, 5% CO_2_, 37°C, and 90% humidity. The cells in the control group were kept in a cell incubator at atmospheric O_2_, 5% CO_2_, 37°C, and 90% humidity. The aCSF was then replaced with Neurobasal Medium supplemented with 100 U/mL penicillin-streptomycin, 1% GlutaMax, and antioxidant-free serum-free B-27 supplement (ThermoFisher Scientific, USA). The cells were maintained in a cell incubator at an atmospheric concentration of O_2_, 5% CO_2_, 37°C, and 90% humidity. The tested compounds CN, ASA, and SC were added either to the incubation culture medium following OGD (reoxygenation) or to both the incubation culture medium and the aCSF at OGD (OGD/reoxygenation) to a final concentration of 0.2 mM, 0.5 mM, or 1 mM. The MTT assay and LDH activity assay were used to assess culture viability 24 hours after OGD.

**Figure 1.**
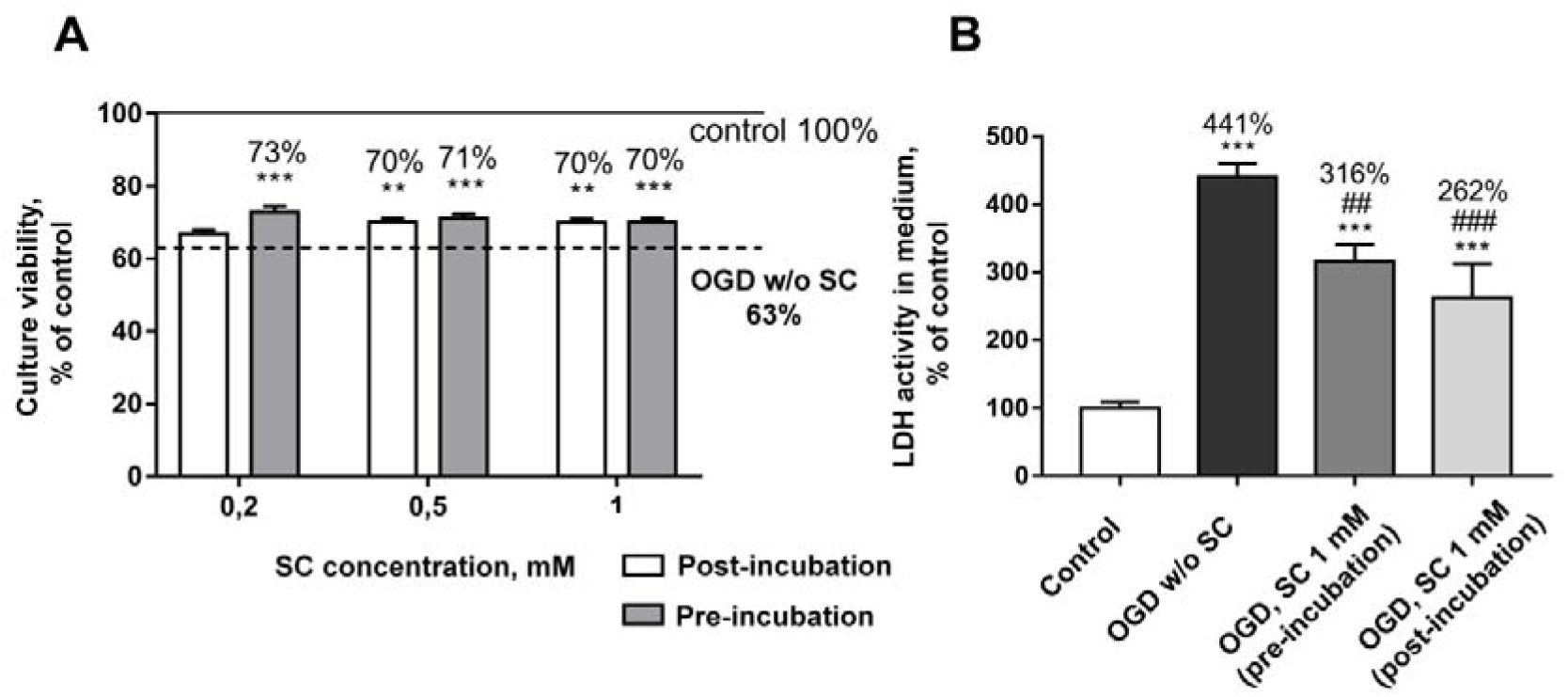
Effect of pre- and post-incubation with 0.2 mM; 0.5 mM, 1 mM SC on the oxygen glucose deprivation (OGD)-induced reduction in viability of primary culture of rat cortical neurons, data are presented as mean ± SEM, N=12, ** - p<0.01; *** - p<0.001 differences from OGD group (dotted line on the histogram) without addition of salicyl-carnosine (SC), viability of culture not subjected to OGD is taken as 100% (solid line on the histogram) (A); Effect of pre-incubation and post-incubation with 1 mM SC on OGD-induced increase in lactate dehydrogenase (LDH) activity in the incubation medium. Data are presented as mean ± SEM, N=8, *** - p<0.001 differences from normoxia group; ## - p<0.01; ### - p<0.001 differences from the OGD group in the absence of SC (B).

### 2.3. NMDA excitotoxicity

We modeled NMDA-induced excitotoxicity according to a previously published method [35]. 10 days following culture seeding, 100 μM glutamate NMDA receptor agonist N-methyl-D-aspartate (NMDA) dissolved in aCSF without magnesium ions, was added to the culture and incubated for 15 min at 37°C, 80% relative humidity, and 5% CO^2^. The current protocol was validated in this study, see Figure 2S. After which all CN, ASA, and SC were added to the incubation medium at final concentrations of 0.2 mM, 0.5 mM, or 1 mM. The MTT assay was used to assess culture viability 24 hours after incubation with NMDA.

**Figure 2.**
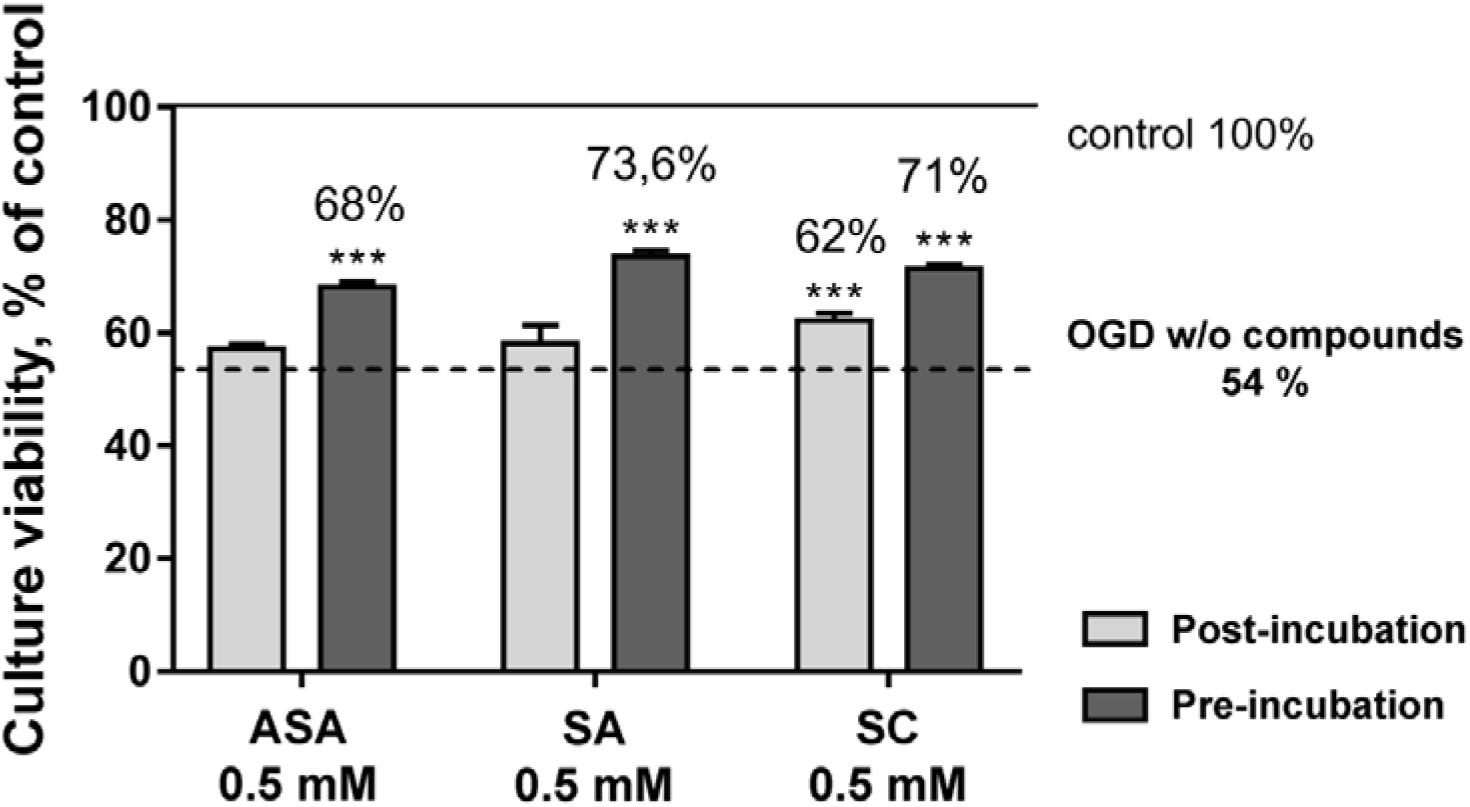
The impact of pre-incubation and post-incubation with 0.5 mM acetylsalicylic acid (ASA), salicylic acid (SA), and salicyl-carnosine (SC) on the decrease in viability of primary rat cortical neurons induced by oxygen glucose deprivation (OGD) (dotted line on the histogram) measured by MTT assay. The viability of the culture that was not subjected to OGD was considered 100% (solid line on the histogram). Data are presented as mean ± SEM, N=12, *** - p<0.001 difference from OGD group without addition of compounds.

### 2.4. MTT assay

Cell viability was assessed in 96-well plates using the MTT assay. The method is based on the reduction of yellow 3-(4,5-dimethyl-2-thiazolyl)-2,5-diphenyl-2H-tetrazolium bromide (MTT) into blue formazan by live cells. The assay was performed as previously described [23].

### 2.5. LDH assay

50 µl of medium was collected into a clean 96-well plate and analyzed according to the protocol of the Lactate Dehydrogenase Activity Assay Kit manufacturer (MAK066, Sigma, USA). The principle of the method consists in the reduction of NAD to NADH by LDH, resulting in the development of color, the intensity of which is determined by colorimetric analysis by the optical density of the solution at a wavelength of 450 nm. Measurement was performed on a Synergy H4 microplate spectrophotometer (BioTek) every 5 min until the value of the most active sample became greater than the value of the NADH standard with the highest concentration (12.5 nmol/well). Values were expressed as the group mean as a percentage of the mean in control wells.

### 2.6. Quantitative determination of salicyl-carnosine in samples

To a 100 μl sample of tissue cell homogenate, 400 μl of a solution of the internal standard (L-alanyl-carnosine, 10 μg/ml) in 10% trichloroacetic acid was added and centrifuged at 16,000 g. The resulting supernatant was injected into the chromatograph loop at a volume of 10 μL.

The analysis was performed using a Finnigan Surveyor LC PumpPlus chromatograph coupled with an LCQ Fleet MS mass spectrometric detector (quadrupole ion trap).

The Ultrasphere 5 ODS analytical column, manufactured by Hichrom Ltd. in the UK, was used for chromatographic separation. The column dimensions were 250×4.6 mm with a 5 μm particle size. The mobile phase consisted of two solutions: 10 mM ammonium acetate acidified with glacial acetic acid to pH 3.7 (solution A) and acetonitrile - 10 mM ammonium acetate (90:10) (solution B) taken in a ratio of 90%A:10%B, and the work was carried out in isocratic elution mode. The flow rate of the mobile phase was 0.7 ml/min, and the sample volume was 10 µl. The separation temperature was 350°C, and the chromatography duration was 10 minutes. The retention time for SC was found to be 5.22±0.05 min, while the retention time for the internal standard (L-alanyl-carnosine) was 4.34±0.05 min. Detection of SC was achieved through mass spectrometry by measuring the daughter product ion with m/z 156.02, which is formed as a result of the decay of the pseudomolecular parent ion with m/z 347.2 at a normalized collision energy of 35 eV. The internal standard (L-alanyl-carnosine) was detected by the total ion current of daughter ions in the range m/z 75 - 300, formed as a result of the decay of the parent ion L-alanyl-carnosine with m/z 298.3.

The mass spectrometer operated in the mode of registering ions positively charged by electrospray (ESI) generated by a voltage of 5 kV. The nebulizer gas (nitrogen) flow rate was 5 l/min, and the nebulizer pressure was 100 psi. The capillary interface temperature was 350°C, and the heater temperature was 300°C. The excitation amplitude at the trap end electrodes was 0.1 V. Helium was used as a damping gas in the ion trap. The data were processed using Xcalibur 2.1 with Foundation 1.0.1 software.

The concentration of SC was measured quantitatively using the internal standard method. The calibration involved measuring the ratio of the chromatographic peak areas of the target substance and the internal standard as a function of the concentration of SC was measured. Calculation was performed by linear regression based on the method of least squares. The calibration dependence was linear within the concentration range of 0.5 µg/mL to 640 µg/mL. The concentration of SC was determined using the formula C = 17.81×S, where C represents the concentration of SC in ng/mL and S represents the area of the chromatographic peak of SC normalized to the area of the internal standard. The methodology for determining SC had a relative error of no more than 10%.

### 2.7. Determination of ROS level in cell culture using DCFH_2_-DA

Primary rat cortical neurons were cultured in 24-well plates. Paraquat was used for inducing oxidative stress. Cells were incubated for 45 minutes after the addition of 100 μM paraquat and the compounds of interest to 4 wells per group. Intracellular ROS formation was detected by measuring fluorescence intensity resulting from staining with DCFH_2_-DA dye. DCFH_2_-DA (10 μM final concentration) was added to the wells 30 minutes before the end of incubation and left in the dark. The cells were then rinsed three times with Hanks’ solution, and the fluorescence level was measured to determine the level of ROS. Fluorescence intensity was detected using a Synergy H1 plate reader (BioTek) and Gen5 software. For this, fluorescence intensity was measured at the bottom of each well using 7×7 points in specific channels (λex 504 nm; λem 524 nm). The average fluorescence intensity was then calculated for each well.

### 2.8. Western blot

Following the experimental procedures, the culture medium was removed. The culture was then washed twice with cold Hanks’ balanced salt solution (PanEco, Russia) and lysed in RIPA-buffer (Sigma, USA), which contained cocktails of protease and phosphatase inhibitors (Sigma, USA). The lysates were centrifuged for 10 minutes at 12000 g, after which the supernatant was aspirated and further analyzed. The protein concentration in the samples was measured using the DC Protein Assay Kit (Bio-Rad, USA). The samples were electrotransferred to a PVDF membrane after Lammlie electrophoresis in a polyacrylamide gel. The membrane was then stained with SYPRO Ruby Protein Gel Stain dye (ThermoFisher, USA) and incubated with primary antibodies to Bak (SC832) and Bax (SC493) (Santa Cruz Biotechnology, USA), Bcl-xL (2764), pERK1/2 (9106), ERK1/2 (4695), p-p38 (4511), p38 (9212), pAkt (4060), Akt (2920), and β-actin (4967) (CellSignaling, USA) and secondary antibodies - anti-rabbit IgG-HRP (7074), anti-mouse IgG-HRP (7076) (Cell Signaling Technology, USA). Immunoreactive bands were detected using the ChemiDoc MP system (Bio-Rad, USA) after visualization with Clarity Max ECL (Bio-Rad, USA).

### 2.9. Statistical processing of data

Statistical data processing was carried out using Microsoft Excel and GraphPad Prism 7 software. The normality of the samples was analyzed using the Shapiro-Wilk test. Multiple comparisons were performed using one-way and two-way ANOVA with Dunnett’s criterion for multiple comparisons.

## RESULTS

### 3.1. Neuroprotective efficacy of various SC concentrations in protecting neurons from oxygen glucose deprivation

Based on the assumptions that SC can exhibit neuroprotective activity in cerebral ischemia, it was decided to investigate its efficacy in the OGD neuronal cell culture model. SC was added to the incubation medium to a final concentration of 0.2 mM, 0.5 mM, or 1 mM either both during OGD and reoxygenation (pre-incubation), or during reoxygenation alone (post-incubation). The neuroprotective efficacy was assessed by evaluating changes in culture viability using the MTT assay. Changes were measured relative to those of the culture subjected to OGD without the addition of SC. Data was analyzed with two-way ANOVA with Tukey’s criterion for multiple comparisons. The results of the MTT assay were confirmed by measuring the change in LDH activity in the incubation medium 24 h after the experiment. Data was analyzed with one-way ANOVA with Dunnett’s criterion for multiple comparisons.

As shown in Figure 1A, culture viability decreases by 37% (p<0.001; t=21.8) following 24 hours of reoxygenation after 4 hours of OGD, compared to the control culture that was not subjected to OGD. During pre-incubation, SC increases culture viability against the decrease induced by OGD at concentrations of 0.2 mM by 10% (p<0.001; t=6.2) and 0.5 mM by 8% (p<0.001; t=5). At concentrations of 0.5 mM and 1 mM, SC increased cell viability by 7% (p<0.001; t=5.2) during OGD and post-incubation (p=0.0015; F=5.9), as measured by the MTT assay (Figure 1A). The neuroprotective effect of SC was confirmed by measuring LDH activity in the incubation medium 24 h after OGD. After 24 hours of exposure to OGD, LDH activity in the culture’s incubation medium increased 4.4-fold compared to the level in the incubation medium of control cells (p<0.001; t=11.9). This indicates an increase in the number of dead cells in the culture, as shown in Figure 1B. Pre-incubation with 1 mM SC resulted in a 1.4-fold decrease in LDH activity (p=0.0017; t=3.8), and post-incubation with 1 mM SC resulted in a 1.7-fold decrease (p=0.001; t=5.1) compared to the culture subjected to OGD without the addition of SC (Figure 1B). These results demonstrate that SC has a neuroprotective effect in conditions of OGD during both pre-incubation and post-incubation of the primary culture of rat cortical neurons in the studied concentration range.

### 3.2. Comparative analysis of the neuroprotective efficacy of 0.5 mM SC, ASA and SA during oxygen glucose deprivation

As SC exerts neuroprotective activity in primary rat cortical neuron culture 24 h after OGD, we compared its effectiveness in this model of cerebral ischemia to that of ASA and SA, which are currently used in the treatment of stroke. During OGD and reoxygenation, SC, ASA, and SA were added to the incubation medium at a final concentration of 0.5 mM. Alternatively, compounds at the same concentration were added during reoxygenation only (post-incubation). The neuroprotective effect was evaluated by measuring the viability of the culture subjected to OGD without any added compounds, using the MTT assay. Data was analyzed with two-way ANOVA with Tukey’s criterion for multiple comparisons.

The primary culture of rat cortical neurons experienced a 46% decrease in viability relative to the control after 4 hours of OGD followed by 24 hours of reoxygenation (Figure 2). The MTT assay showed that all three compounds at a concentration of 0.5 mM increased culture viability during pre-incubation compared to the culture subjected to OGD without the addition of these compounds (p<0.001, F=163). ASA, SA, and SC were found to increase viability by 14% (p<0.001, t=12.7), 19.4% (p<0.001, t=17.6), and 17% (p<0.001, t=17.6), respectively (Figure 2). Only SC was found to be neuroprotective when added during reoxygenation, resulting in an 8% (p<0.001, t=4.2) increase in culture viability compared to cultures that were subjected to OGD and not incubated with the tested compounds (Figure 2).

### 3.3. Comparative analysis of the neuroprotective effect of exposure to 0.5 mM SC, ASA and CN for 24 h after 15 min of exposure to 100 µM NMDA

It was investigated whether SC can provide neuroprotection against NMDA-induced excitotoxicity in primary rat cortical neuron culture, which is one of the main causes of neuronal death in ischemic brain damage. CN and ASA were used as drugs for comparison. After incubation with 100 µM NMDA for 15 minutes, all three substances were added to the incubation medium at final concentrations of 0.2 mM, 0.5 mM, or 1 mM. The MTT assay was used to assess culture viability 24 hours after incubation with NMDA. Data was analyzed with one-way ANOVA with Dunnett’s criterion for multiple comparisons.

As shown in Figure 3A, 15-minute incubation with 100 µM NMDA resulted in a 26% decrease in viability (p<0.001; F=50.2) of the primary rat cortical neuron culture 24 hours post-incubation. The addition of 0.5 mM and 1 mM SC caused a 6% (p<0.001; F=15.9) and 12% (p<0.001; t=6.5) increase in viability (p<0.001; F=15.9) of the culture pre-incubated with NMDA, respectively (Figure 3A). Pre-incubation with NMDA followed by incubation with 1 mM CN resulted in an 8% increase in culture viability (p<0.001; F=11.7) compared to the control (Figure 3B). Similarly, addition of 1 mM ASA resulted in an 11% increase in viability (p<0.001; F=8.7) of the culture pre-incubated with NMDA (Figure 3B). Therefore, SC has the most pronounced neuroprotective effect, even at concentration 2 times lower than CN and ASA (0.5 mM) following acute exposure to NMDA.

**Figure 3.**
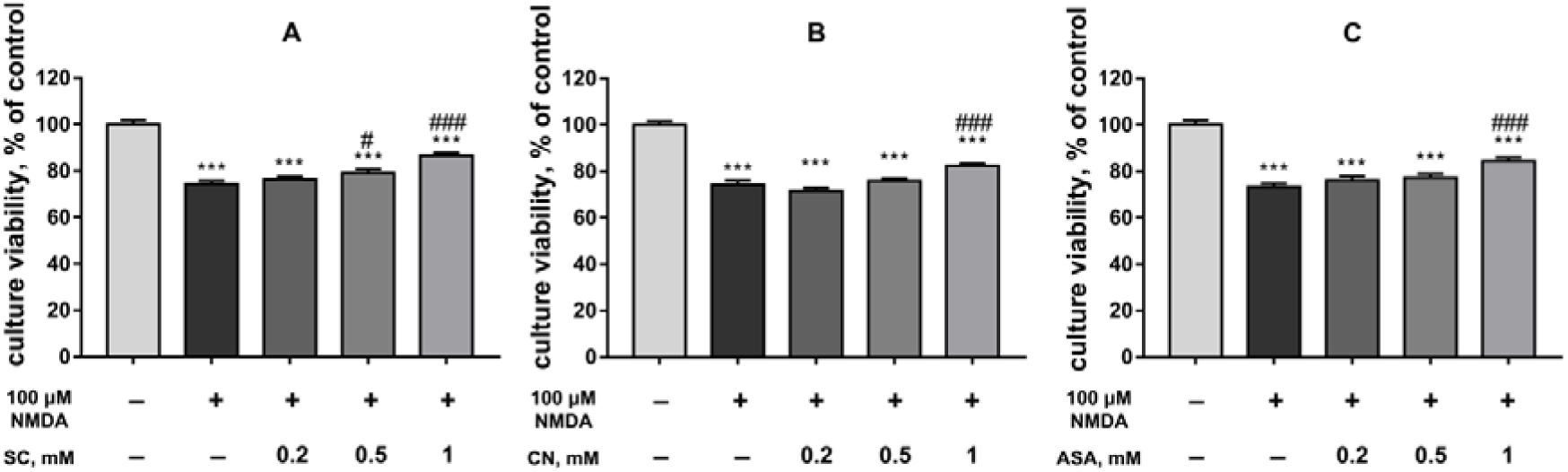
Effect of 24 h incubation of primary rat cortical neuron culture with 0.2 mM, 0.5 mM, and 1 mM salicyl-carnosine (SC) (A), carnosine (CN) (B) and acetylsalicylic acid (ASA) (C) on the decrease in viability induced by 15 min pre-incubation with 100 µM NMDA. Data are presented as mean ± SEM, N=12, *** - p<0.001 difference from control; # - p<0.05, ### - p<0.001, differences from culture incubated with NMDA without the addition of compounds of interest.

### 3.4. SC uptake by primary culture rat cortical neurons

The question arose as to the mechanism of the neuroprotective effect of SC, since SC exhibited pronounced neuroprotective activity in models of ischemia on primary rat cortical cell culture. A direct antioxidative effect is most likely, since SC has previously been shown to have antioxidative activity *in vitro* [32]. However, in order for this mechanism to be viable, SC must pass through the cytoplasmic membrane. To address this question, we determined whether SC can penetrate neurons effectively. Primary rat cortical neuron cultures were incubated with 2 mM and 20 mM SC dissolved in aCSF for 30 minutes. The cells were then washed, lysed, and the amount of SC present in the lysate was evaluated.

SC can effectively pass into neurons in proportion to its concentration in the sample. After incubating the culture with 2 mM SC for 30 minutes, the amount of SC in the lysate was 1.36±0.12 nmol/mg protein (N=3). When the culture was incubated with 20 mM SC, the amount was 9.95±0.79 nmol/mg protein (N=3). Due to its ability to pass into neurons, it can be concluded that SC is capable of exerting a neuroprotective effect in models of ischemia by acting on targets within the cytoplasm, including the possibility of direct antioxidant action.

### 3.5. SC increases culture viability under conditions of paraquat-induced OS as assessed by the MTT assay

To determine whether SC can inhibit the development of OS in neurons, the neuroprotective efficacy of SC was evaluated on primary cultures of rat cortical neurons under conditions of paraquat-induced OS. The cultures were incubated with 100 μM paraquat for 48 hours, with or without the addition of 0.2 mM, 0.5 mM or 1 mM SC to the incubation medium. Culture viability was assessed using the MTT assay. Data was analyzed with one-way ANOVA with Dunnett’s criterion for multiple comparisons.

As shown in Figure 4, the viability of the culture was reduced by 77.4±1.2% compared to intact cells following 48 hours of incubation with 100 µM paraquat. The addition of 1 mM SC to the incubation medium increased culture viability by 19±5.2% compared to the 100 μM paraquat group. However, the addition of 0.2 mM and 0.5 mM SC to the incubation medium did not significantly increase culture viability when exposed to 100 μM paraquat. Therefore, in the model of OS induced by 48-hour incubation with 100 μM paraquat, SC at a concentration of 1 mM showed neuroprotective properties.

**Figure 4.**
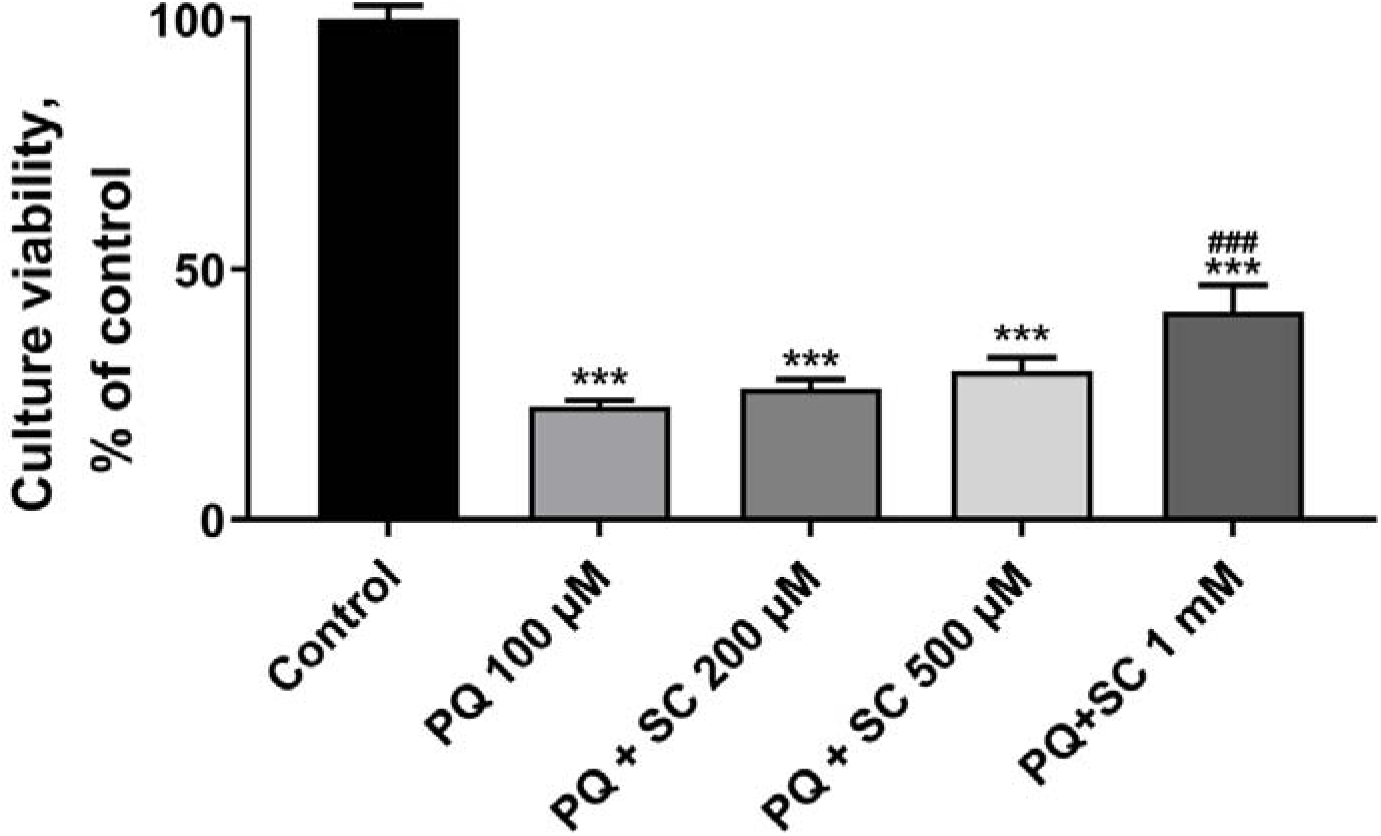
Effect of 200 μM, 500 μM, 1 mM SC on the viability of primary rat cortical neuron culture when incubated with 100 μM paraquat for 48 h according to the MTT assay. Data are presented as mean ± SEM, N=12; *** - p<0.001 difference from control group; ### - p<0.001 difference from 100 μM paraquat group.

### 3.6. Effect of SC on ROS levels under conditions of paraquat-induced OS

The impact of SC on ROS levels was assessed in primary rat cortical neuron cultures in conditions of paraquat-induced OS. DCF fluorescence intensity, which indicates intracellular ROS levels, was measured by adding 1 mM SC to the incubation medium during exposure to 100 μM paraquat. Data was analyzed with two-way ANOVA with Tukey’s criterion for multiple comparisons.

As shown in Figure 5, a 45-minute incubation of neurons with 100 μM paraquat increased DCF fluorescence intensity by 35.7±4.5% (p<0.001) compared to intact cells. After 45 minutes of incubation, 1 mM SC caused a 47.2±11% (p<0.001) decrease in DCF fluorescence intensity relative to cells incubated with 100 μM paraquat. The DCF fluorescence intensity in cultures incubated for 45 minutes with 1 mM SC alone did not differ from the DCF fluorescence intensity in intact cultures. Thus, the paraquat-induced increase in ROS in a primary culture of rat cortical neurons is prevented by the inclusion of 1 mM SC in the incubation medium.

**Figure 5.**
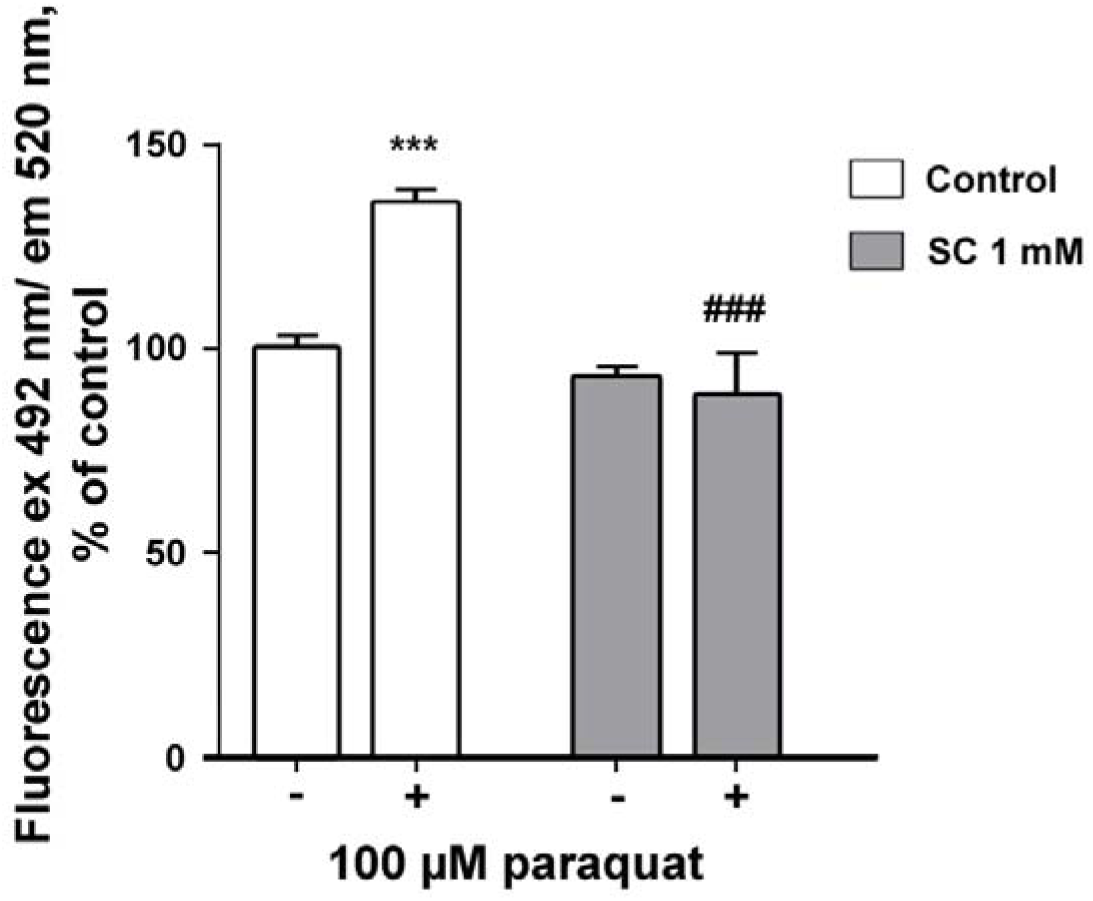
The effect of 1 mM SC on DCF fluorescence intensity in primary rat cortical neuron culture induced by 45 min incubation with 100 μM paraquat. The Data are expressed as % of the mean ± SEM of the DCF fluorescence intensity (excitation at 492 nm wavelength, emission at 520 nm) in the control wells of the plate, N=8; *** - p<0.001 - difference from intact cells; ### - p<0.001 - difference from culture incubated with 100 μM paraquat.

### 3.7. Impact of SC on signaling cascades associated with neural apoptosis during paraquat-induced OS

The antioxidative activity of SC was expected to reduce the activation of intracellular processes related to OS and neuron death within the culture cells in the context of paraquat-induced OS. To test this hypothesis, we compared the levels of proapoptotic and antiapoptotic proteins from the Bcl-2 family in lysates of cultures incubated for 3 hours with 100 μM paraquat or 100 μM paraquat and 1 mM SC, using Western blotting. Data was analyzed with two-way ANOVA with Tukey’s criterion for multiple comparisons.

As shown in Figure 6A, incubation of primary rat cortical neuron cultures with 100 µM paraquat for 3 hours resulted in an 88% increase in Bak levels in the culture relative to control (p = 0.0127). However, when the culture was incubated with 100 μM paraquat in the presence of 1 mM SC, the amount of Bak did not increase relative to control and was 73% lower than when the culture was incubated with 100 μM paraquat alone (p = 0.0457). Bax and Bcl-xL levels were not significantly altered in all groups, including the control (Figure 6B, C). Thus, 1 mM SC reduces the paraquat-induced increase in the amount of the pro-apoptotic protein Bak in primary culture neurons, while having no significant effect on Bax and Bcl-XL.

**Figure 6.**
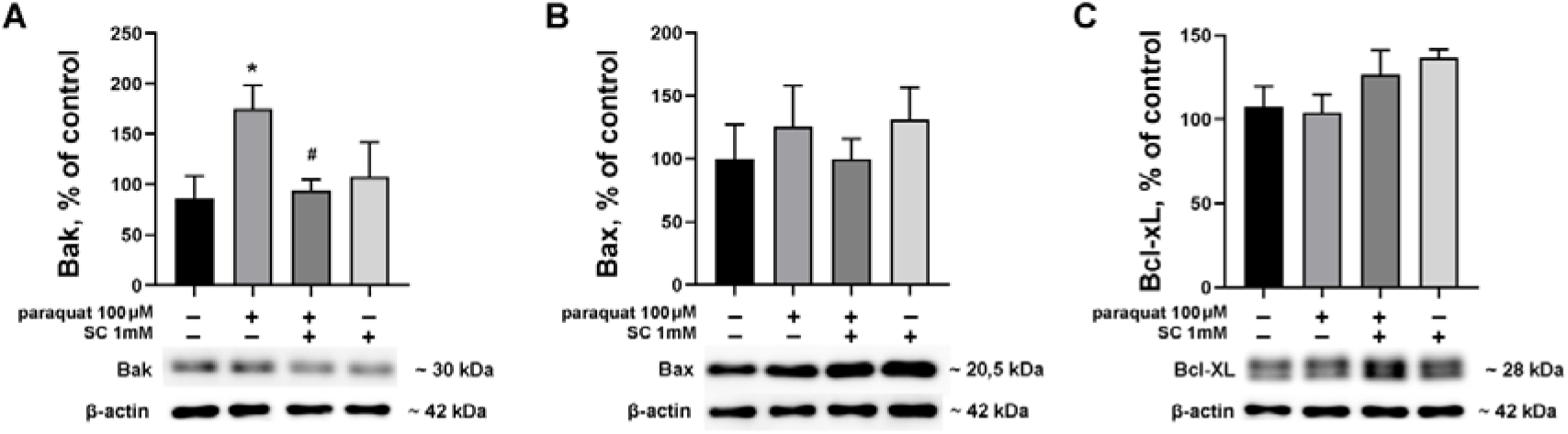
Effect of 3 hour incubation of primary rat cortical neuron culture with 100 μM paraquat alone and in the presence of 1 mM SC on the amount of Bak (A), Bax (B) and Bcl-XL (C) proteins. Data are presented as % of control ± SEM, N=6; * - p < 0.05, # - p < 0.05. Representative images of immunoreactive bands are shown below the graphs.

We also assessed the activation of intracellular signaling cascades associated with OS and the regulation of neuronal viability in culture lysates incubated for 3 h with 100 μM paraquat or 100 μM paraquat and 1 mM SC. The ratio of phosphorylated (activated) and total forms of ERK1/2, p38, and Akt kinases were compared among all experimental groups.

Figure 7 presents changes in the activation levels of the ERK 1/2, p38 and Akt kinases after a 3-hour incubation of a primary culture of rat cerebral cortex neurons with 100 μM paraquat, 1 mM SC and 100 μM paraquat in the presence of 1 mM SC. The experiment revealed no significant difference in the activation of ERK 1/2 kinase in culture following incubation in the presence of paraquat compared to the control. However, following a 3-hour incubation with 100 μM paraquat in the presence of 1mM SC, ERK 1/2 activation increased by 172% compared to the control (p = 0.0234). Additionally, a significant difference in ERK 1/2 activation was observed between the 100 μM paraquat + SC group and the 1 μM SC group (85% and 273% of control, respectively, p = 0.0180 (Figure 7A)).

**Figure 7.**
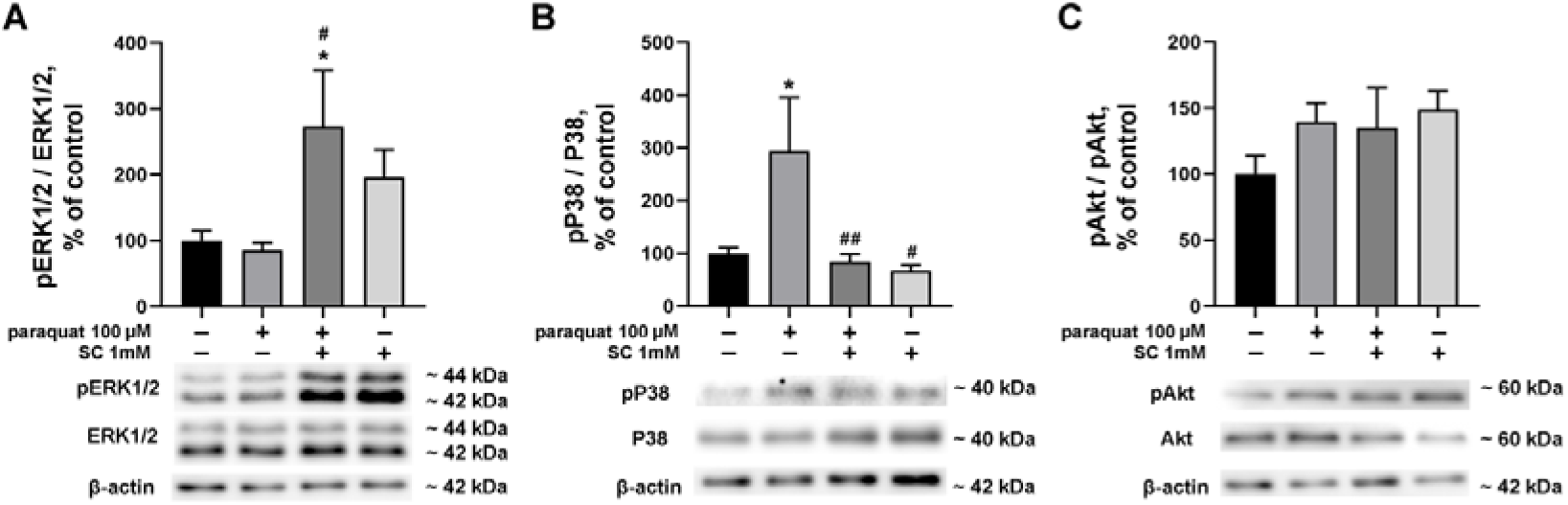
Effect of 3 hour incubation of a primary culture of rat cortical neurons with 100 μM paraquat alone and in the presence of 1 mM SC on the activation of ERK 1/2 (A), p38 (B), and Akt (C) kinases. The data are presented as a percentage of control ± SEM, N=6; * - p < 0.05, # - p < 0.05, ## - p < 0.01. Representative images of immunoreactive bands are shown below the graphs.

After a 3-hour incubation with 100 μM paraquat, p38 kinase activation increased by 194% compared to the control (p = 0.0171) (Figure 7B). The addition of 1 mM SC to the medium during culture incubation with 100 μM paraquat resulted in a decrease in p38 activation compared to the 100 μM paraquat group, bringing it down to the control level (p = 0.0098 relative to the 100 μM paraquat group). No significant differences in p38 activation were found between the 1 mM SC and control groups. However, there was a difference in p38 activation between the 100 μM paraquat and 1 mM SC groups, with the former showing 294% and the latter showing 67% of control.

There were no significant differences in Akt kinase activation observed among the studied groups (Figure 7C). It can therefore be concluded that 1 mM SC reduces the paraquat-induced increase in p38 kinase activation without significantly affecting Akt kinase activation. Additionally, when co-incubated with 100 μM paraquat, 1 mM SC leads to increased activation of ERK 1/2 kinase.

## DISCUSSION

A common method for modeling neural tissue ischemia is oxygen glucose deprivation (OGD) of neuronal cell cultures. Depending on the type of cultured cells, the culture maintenance conditions and the resources available in a given laboratory, the duration of OGD and the percentage of cell death can vary. In this study, a primary culture of rat cortical cells was subjected to 4 hours of OGD, resulting in a decrease in culture viability of between 37% and 46%. However, this model is the most suitable for reproducing both the ischemic phase, including the release of excitotoxic neurotransmitters [36], and the reoxygenation phase in neuronal cultures, thereby being suitable for studying the processes occurring in both phases, as well as for screening neuroprotective compounds and evaluating their neuroprotective efficacy[33, 37–39].

According to the obtained data, SC at concentrations ranging from 200 µM to 1 mM reduces neuronal death by one third in a culture model of OGD. This efficacy is either comparable or superior to other compounds active in this model, such as resveratrol [32], oxycontin [35], ginsenoside Rd [36], and L-carnitine [37]. Although the neuroprotective efficacy of SC when added at the beginning of OGD is comparable to SA and ASA, only SC shows a neuroprotective effect when added during reoxygenation. This fact gives SC an advantage over the comparison drugs (ASA and SA) because, as mentioned above, the use of neuroprotective agents during the reperfusion and recovery phase can significantly improve the patient’s rehabilitation prognosis.

As previously stated, excitotoxicity is one of the mechanisms of neuronal death that occurs immediately after the onset of cerebral ischemia [40]. The activation of extrasynaptic NMDA receptors containing NR2B by glutamate is crucial in the development of excitotoxicity [41]. Prolonged activation of NMDA receptors leads to the influx of excess Ca^2+^ into the cell, resulting in mitochondrial damage, and initiating the mitochondrial apoptosis pathway and OS [42]. This study demonstrates that SC can effectively prevent NMDA-induced cell death in primary cultures of rat cortical neurons. Additionally, SC exhibits neuroprotective properties in NMDA-induced excitotoxicity at a lower concentration than the comparison drugs, ASA and CN.

In order for a compound to have a direct antioxidant effect, it must be capable of entering neurons. SC is taken up by primary cultured rat cortical neurons at concentrations that are neuroprotective in the models investigated. The mechanism of its uptake is beyond the scope of this study and is subject to further investigation. However, the ability to enter neurons makes SC a promising compound for neutralizing ROS generated in mitochondria during reoxygenation [8].

Previous research by our group has demonstrated that SC exhibits direct antioxidant activity in models of iron-induced chemiluminescence in serum [32]. The current study shows that SC reduces paraquat-induced neuronal death, as measured by the MTT assay, and decreases the amount of ROS detected by DCF fluorescence. Paraquat is a herbicide commonly used to model various pathologies *in vitro*. Its toxicity to cells is caused by an increase in intracellular reactive oxygen species (ROS), leading to oxidative stress (OS) and triggering pro-apoptotic signaling cascades [43]. SC’s efficacy in protecting neurons from paraquat-induced OS suggests that it may directly prevent OS development.

A neuroprotective compound should be able to prevent the triggering of proapoptotic signaling cascades induced by OS. In a model of paraquat-induced OS, addition of SC prevented an increase in levels of proapoptotic protein Bak in primary rat cortical culture. In the ischemic core, neuronal loss is predominantly necrotic; in the penumbra, neurons have diminished function and are lost predominantly via the mitochondrial pathway of apoptosis [44]. Although both Bak and Bax expression is upregulated in tissue during cerebral ischemia [45], paraquat exposure causes increased Bak, but not Bax, expression. Bak knockdown reduces paraquat toxicity [46]. Thus, in this study we have shown that the neuroprotective activity of SC during OS induction is associated with decreased expression of the proapoptotic protein Bak, which contributes to the activation of the mitochondrial apoptotic pathway.

Activation of a number of intracellular signaling cascades mediates the response of neurons to excitotoxic neurotransmitter release and OS. We have shown that SC protects against paraquat-induced OS by inhibiting the activation of MAP kinase p38. Previously, p38 activation has been shown to mediate neuronal apoptosis upon exposure to glutamate or NMDA and OS induction by mitochondrial toxins [47–49]. A number of signaling cascades associated with neuronal protection against OS are also activated by cerebral ischemia. The activation of the key signaling pathways Raf1/ERK and PI3K/Akt is associated with the survival of neurons in the penumbra [50]. The SC-induced increase in ERK1/2 activation may also contribute to its neuroprotective effect in the paraquat-induced OS model. Thus, both a direct antioxidant effect within neurons and the prevention of proapoptotic signaling cascades and the stimulation of antiapoptotic signaling cascades are involved in the mechanism of the neuroprotective effect of SC.

## CONCLUSIONS

This study demonstrated high neuroprotective efficacy of SC in modeling the main pathogenetic factors causing neuronal death in ischemic stroke on primary rat cortical cell culture: excitotoxicity and OS. It has been shown that SC is able to effectively penetrate inside neurons and prevent the development of OS by reducing the amount of ROS inside neurons. Also, the neuroprotective properties of SC are associated with suppression of activation of pro-apoptotic signaling cascades inside neurons. These *in vitro* effects of SC make it a promising compound for further studies in models of cerebral ischemia.

## Author contributions

Conceptualization: Fedorova Tatiana N., Lopachev Alexander V.; Methodology: Lopachev Alexander V., Kazanskaya Rogneda B., Kulikova Olga I., Abaimov Denis A.; Formal analysis and investigation: Lopachev Alexander V., Kazanskaya Rogneda B., Kulikova Olga I., Khutorova Anastasiya V., Abaimov Denis A.; Writing - original draft preparation: Lopachev Alexander V., Kazanskaya Rogneda B.; Writing - review and editing: Kulikova Olga I., Khutorova Anastasiya V., Abaimov Denis A., Fedorova Tatiana N.; Funding acquisition: Fedorova Tatiana N.; Resources: Fedorova Tatiana N., Abaimov Denis A.; Supervision: Fedorova Tatiana N.

## Acknowledgments

This work was supported by state assignment Ministry of Education and Science of the Russian Federation for Research Center of Neurology № АААА-А20-120052790037-4.

## Ethics declarations

The authors have no competing interests to declare that are relevant to the content of this article. Animal Ethics declaration: not applicable

## Abbreviations

SC: salicyl-carnosine
NMDA: N-methyl-D-aspartate
ERK1/2: Extracellular Signal-Regulated Kinase 1/2
ATP: adenosine triphosphate
AMPA: α-amino-3-hydroxy-5-methyl-4-isoxazolepropionic acid
ROS: reactive oxygen species
OS: oxidative stress
OGD: glucose-oxygen deprivation
ASA: acetylsalicylic acid
CN: carnosine
PEPT2: peptide transporter 2
EDTA: ethylenediaminetetraacetic acid
FBS: fetal bovine serum
aCSF: artificial cerebrospinal fluid
LDH: lactate dehydrogenase
NAD: nicotinamide adenine dinucleotide
DCFH_2_-DA: dichlorodihydrofluorescein diacetate
PVDF: polyvinylidene fluoride
Bcl-2: B-cell lymphoma 2 protein
Bax: Bcl-2 Associated X protein
Bak: Bcl-2 homologous antagonist/killer protein
Bcl-xL: B-cell lymphoma-extra large protein
MAP kinase: mitogen-activated protein kinase
Akt: protein kinase B
PI3K: phosphoinositide 3-kinase.

